# The Effects of Hypothermic Storage on Passaged Chondrocyte Viability and Redifferentiation Potential

**DOI:** 10.1101/2025.06.20.660742

**Authors:** Dilpreet Rayat, Jack Miller, Stephanie Richardson-Solorazano, Rylee E. King, Valerie C. West, Alvin W. Su, Justin Parreno

**Author notes:** Equal Contributions (Dilpreet Rayat, Jack Miller).

## Abstract

Cell-based transplantation therapies, such as autologous chondrocyte implantation (ACI), are used to treat focal cartilage defects caused by trauma or degeneration. In ACI, chondrocytes are isolated from non-load-bearing regions of healthy cartilage regions and then sent to a cell manufacturing laboratory, where they are expanded for cell number in monolayer culture. Once a large number of cells are obtained, they are transported to the clinic for reimplantation into the defect site. The storage and transport conditions from cell manufacturing to implantation may be a critical time that could influence cell viability and redifferentiation potential.

Although hypothermic storage at sub-physiological temperatures is commonly used to preserve cell viability, long-term storage of cartilage under hypothermic conditions can impair chondrocyte viability and function. However, the impact of short-term, acute hypothermic storage on passaged chondrocytes remains largely unknown. We tested the hypothesis that acute hypothermic storage negatively impacts passaged chondrocyte viability and reduces the capacity for redifferentiation.

Passaged chondrocytes were stored either in monolayer culture or in suspension at 36, 19 or 8°C. In monolayer culture, hypothermic temperatures preserved cell viability with no difference compared to storage at 36°C for up to three days. Additionally, hypothermic temperatures promoted cell rounding, reduced proliferative capacity, depolymerized filamentous actin, and led to a slight reduction in the mRNA levels of specific matrix molecules compared to 36°C. Intriguingly, the effects of hypothermia were context-dependent. Exposure of passaged cells in suspension to hypothermia promoted the maintenance of cell viability and reduced aggregation compared to 36°C. When stored at 8°C in suspension, passaged cells exhibited enhanced expression of specific matrix molecule mRNA levels compared to cells at 36°C in suspension. Subsequently, when passaged cells in suspension at 8°C were seeded in 3D within adherent agarose molds, there was an increase in aggrecan expression 10 days after seeding. The tissues formed by cells stored in suspension at 9°C were thicker and stained more intensely for aggrecan. Therefore, in contrast to our hypothesis, we found that hypothermic storage did not have a negative impact; when stored for 1 day in suspension, it had lasting effects on matrix deposition. The storage of passaged chondrocytes under hypothermic conditions may be beneficial for ACI, warranting further investigations of cell hypothermic storage for *in vivo* repair.

## Introduction

Autologous chondrocyte implantation (ACI) is a cell transplantation therapy used to treat focal cartilage damage[1–3]. In ACI, during an initial arthroscopy procedure, healthy cartilage is harvested from a non-load-bearing region of the patient’s articular cartilage and shipped to a cell manufacturing laboratory. In the laboratory, chondrocytes are isolated from harvested cartilage pieces and then expanded to increase cell number in monolayer, two-dimensional (2D) culture. Once a large number of cells are acquired, the resultant cells are transported from the cell manufacturing laboratory to the clinic for surgical implantation. The success of ACI is in part dependent on the transplantation of viable cells capable of producing cartilaginous repair tissue [4].

The cellular environment in which cells are maintained is a major determinant of viability and cartilage matrix production. While culturing chondrocytes in monolayer on stiff polystyrene enables the proliferation for the generation of a large number of cells, monolayer expansion also leads to chondrocyte dedifferentiation. In dedifferentiation, chondrocytes alter in morphology by flattening and increasing in area [5]. In addition, chondrocytes alter their cytoskeleton by reorganizing filamentous (F-) actin from a cortical to a stress fiber organization [5–11]. The altered morphology and cytoskeleton lead to dysregulation of matrix molecule expression. Rather than expressing cartilage matrix molecules, such as type II collagen (Col2) and aggrecan (Acan), monolayer-expanded (passaged) chondrocytes express fibrous matrix molecules, including type I collagen (Col1) and Tenascin C (Tnc). This results in the production of fibrocartilage, which is biomechanically inferior to articular cartilage [8, 12]. Furthermore, passaged chondrocytes also produce contractile molecules, such as alpha-smooth muscle actin (αSMA) and transgelin (TAGLN) [13–15]. The expression of contractile molecules enhances the propensity for tissue contraction which may affect tissue integration. Chondrocyte dedifferentiation, resulting from the monolayer culture environment, may compromise the outcomes of ACI.

In addition to monolayer culture, hypothermic storage of chondrocytes is an environmental variable that can impact cellular health and matrix-producing capabilities. During monolayer expansion, cells are maintained in standard cell culture incubators set to physiological body temperature (∼37°C). However, following cell expansion, cells are transported from the cell manufacturing laboratory back to the clinic, often in suspension culture, at temperatures below physiological temperature. There remains limited knowledge on the effect of acute, short-term storage at sub-physiological (hypothermic) temperatures on the viability, phenotype, and ability to form repair matrix by passaged chondrocytes. However, in general, the culture of chondrocytes under hypothermic conditions is regarded as unfavorable. Storing goat osteochondral tissues at 4°C [16] or human cartilage discs at 22°C [17] reduces chondrocyte viability as compared to storage at 37°C. Culturing nasal chondrocytes at 32.2°C slows proliferation [18]. Culture of primary chondrocytes at 32°C reduces glycosaminoglycan accumulation in pellet culture [19]. Furthermore, prolonged culturing of passaged chondrocytes at 32°C impairs the redifferentiation of passaged chondrocytes [20]. This is consistent with studies in other cell types, where cooler temperatures also compromise cellular viability and health of non-chondrocyte mammalian cells [21].

Since there is limited knowledge on the effect of acute, short-term hypothermic temperatures on passaged chondrocytes specifically, we sought to test the hypothesis that acute storage of passaged chondrocytes at hypothermic temperatures will be detrimental, causing an increase in cellular death and reducing the chondrogenic phenotype and capacity to produce matrix by passaged cells.

## Materials and Methods

### Chondrocyte isolation and expansion

Bovine metacarpal phalangeal joints were procured from a local butcher. Under aseptic conditions, full thickness articular cartilage were scraped from the joints using a scalpel. Chondrocytes were liberated from tissues by matrix digestion in a CO2 incubator first using 0.5% proteinase Type XIV derived from Streptomyces griseus (P5147; Sigma-Aldrich; Burlington, MA, USA) for 1 hour, followed by 0.1% collagenase (Roche; Brulington, MA, USA) digestion overnight. Digests were filtered through a 100μm cell strainer and pelleted by centrifugation at 600g. Cells were washed twice in serum-free Ham’s F12 (Corning; Manassas, VA, USA) and then resuspended in growth ‘complete media’ which consisted of Ham’s F12 supplemented with 10%FBS (GenClone; Genesee Scientific, San Diego, CA, USA) and antimycotic-antibiotic (MilliporeSigma; Burlington, MA, USA). Cells were expanded in monolayer by seeding freshly isolated (passage 0; P0) cells at density of 2 x 10^3^ cells/cm^2^ on polystyrene T-flasks (Cat#25-111; GenClone). Cell expansion was performed by maintaining cells in a CO2 incubator. Media was replenished every 3 days with fresh complete media. Once cells reached 70-90% confluency, cells were removed from flasks using 0.25% Trypsin/EDTA (GenClone). At this point cells were deemed as passage 1 (P1). Cells were reseeded onto T-flasks at a density 2 x 10^3^ cells/cm2 for further expansion to passage 2 (P2).

### Cell culture under hypothermic conditions

Once P2 cells reached >80% confluency, they were removed from flasks. Cells were pelleted, resuspended in fresh complete media. To investigate the effects of temperature on adherent P2 cells, cells were seeded at a density of 2.5 x 10^5^ cells per cm^2^ onto either wells of a 6-well dish or onto glass dishes (FD35-100; WPI; Sarasota, FL, USA) for immunofluorescent imaging. Cells were maintained in 2mL of complete media in a temperature-controlled CO2 incubator. After 24 hours, cells were placed under hypothermic conditions by placing into a Styrofoam container and maintained under ambient (room temperature) conditions on a laboratory bench, or placing cells in a temperature-controlled merchandiser refrigerator. Control cells were maintained in the temperature-controlled CO2 incubator. To investigate the effects of temperature on P2 cells in suspension, P2 cells from tissue culture flasks were pelleted and resuspended in fresh complete media at a density of 1 x 10^6^ cells per mL and maintained in 15mL conical tubes. The temperatures of the various environmental conditions were monitored using high accuracy temperature data loggers (±0.3°C accuracy) (iDiAK; Amazon).

### 3D redifferentiation culture of passaged chondrocytes

3D cultures were prepared in agarose molds as previously described with slight modifications [22–24]. Briefly, agarose molds created by pipetting 2mL of molten agarose within wells of a 12-well plate (Falcon; Fisher Scientific, Waltham, MA, USA). Following agarose gelation, 8mm biopsy punches were used to core agarose at the center of wells. Molds were washed using PBS (Quality Biological, Gaithersburg, MD, USA), prior to seeding of cells within molds.

To investigate the effect of hypothermic temperatures on the matrix forming capabilities of passaged cells, once P2 cells reached >90% confluency, they were removed from flasks using 0.25% Trypsin and resuspend in complete media at a density of 1 x 10^6^ cells per mL. Cell suspensions were maintained at the various environmental temperatures as described above. After 24 hours, cells were prepared for 3D culture. Suspended cells were pelleted by centrifugation and resuspended in fresh complete media at a density of 1.0 x 10^7^ cells per mL. Approximately, 2 x 10^6^ cells were seeded within each mold by pipetting 200uL of cell suspension into molds. Cells were left undisturbed to allow for initial attachment to underlying polystyrene. After 3 hours, 1.5mL of fresh complete media was gently placed into wells. Cells seeded in 2D polystyrene as well as that were seeded for 3D culture immediately following detachment from 2D culture flasks served as experimental controls. Media consisting of DMEM containing 1x antimycotic/antibiotic (Corning, Edison, New Jersey, USA), 40ug/mL L-Proline (Sigma Aldrich, St. Louis, MO, USA), 0.1um Dexamethasone (Sigma Aldrich, St. Louis, MO, USA) and 100ug/mL L-ascorbic acid (Sigma Aldrich, St. Louis, MO, USA)), which we define as ‘redifferentiation media’, was used to stimulate redifferentiation. Redifferentiation media was introduced into cultures in a mix with complete media, beginning on day 2. In addition, media was supplemented, at various time points listed in Table S1, with 10uM Y27632 (Selleckchem, Houston, TX, USA) and 10ng/mL TGFβ3 (R&D Systems, Minneapolis, MN, USA), to prevent matrix contraction and enhance matrix deposition, respectively.

### Cell Morphology

Cell morphology of P2 chondrocytes in 2D culture was quantified from light microscope images captured using a Swift microscope digital camera mounted on a Zeiss microscope. Image quantification was performed using FIJI ImageJ as previously described. To measure the ability of passaged chondrocytes to recover after hypothermic storage, adhered cells were stored in hypothermic conditions (19 or 8°C) for 24 hours and then allowed to recover at 36 °C. Following hypothermic storage, the cells were placed in a CO2 incubator for 48 hours. Recovery was assessed by examining changes in cell morphology using light microscope images captured, 24 and 48 hours later, with a Swift digital microscope camera mounted on a Zeiss microscope. Cell area and circularity were used to quantify cell morphology and were measured in FIJI ImageJ as previously described [14, 25]. In comparison to their morphology immediately after hypothermic storage, the cells showed partial to full re-spreading. There were also increases in cell area, as well as a decrease in circularity, suggesting that cells can recover their morphology following hypothermic storage.

Cell area and circularity were quantified as previously described [25]. Briefly, cell boundaries were manually traced using light microscopy images on FIJI ImageJ software. Cell area and circularity were measured, where circularity was defined as C = 4π(A/P^2^) where P is the perimeter of the cell and A is the area of the cell. Circularity values of 0 represent elongated ellipses and values of 1 represent perfect circles.

### Chondrocyte viability

Adhered cells cultured on glass were transferred into fresh complete media containing Viaflour-CF488 (1:1000; Biotium; Fremont, CA, USA), Live-or-Dye 568 (1:1000; Biotium), and Hoechst (1:500; Biotium). Cells were stained in a CO2 incubator. After 3 minutes, cells were rinsed in PBS three times and then fixed using 4% paraformaldehyde (PFA). After 15 minutes of fixation, cells were washed in PBS twice, and then coverslip mounted using Everbrite mounting medium (Biotium).

Cells in suspension were pelleted by centrifugation at 600g and resuspended in fresh complete media that contained Viafluor (1:1000), Live-or-Dye (1:1000) or Live-or-Dye 568 (1:1000), and Hoechst 33342 (1:500) to preferentially stain for live cells, dead cells, or all cell nuclei. Cells were stained for 30 minutes in a CO2 incubator. Cells were pelleted by centrifugation, and resuspension in 1mL PBS to wash cells. After two washes in PBS, cells were fixed in 4% PFA. After 15 minutes, cells were washed in PBS to remove fixative and resuspended in PBS. Immediately prior to imaging, 20uL of cell suspension was placed in the center of a glass dish and an 18mm glass coverslip was placed on cells to restrict motility within the dish during imaging. Cells were imaged on a Zeiss LSM880 confocal microscope using a 20x objective.

Cell viability was quantified using the formula 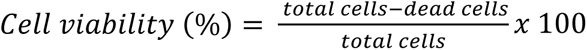, where total cells are the cells stained for Hoechst and dead cells are cells positive for Live-or-Dye and low staining for Viafluor. To calculate cell density, the formula 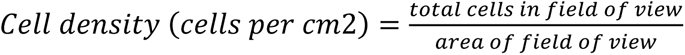.

### Actin staining and Quantification

Cells on glass dishes were fixed in 4%PFA for 15 minutes. Following fixation, cells were permeabilized for 30 minutes in permeabilization/blocking buffer (3% Goat serum, 3% BSA, and 0.3% Triton in PBS). Cells were then incubated in staining solution containing Rhodamine phalloidin (1:50; Biotium) to stain F-actin, Vitamin D-binding protein conjugated to AlexaFluor488 (1:200; RayBiotech; Peachtree Corners, GA, USA) to stain G-actin, and Hoechst (1:500) to stain nuclei. After 1 hour at room temperature, cells were washed in six washes of PBS, and then glass coverslipped using Everbrite mounting medium. Cells were imaged on a Zeiss LSM880 confocal microscope using a 40x objective.

To quantify the proportion of cells displaying stress fibers, confocal images were anonymized and provided to an individual blinded to the experimental conditions. Using FIJI Image J software, the total cell number in each image was quantified by counting the number of nuclei. Next, cells were evaluated for the presence of stress fibers, which are arrays of parallel F-actin bundles. Cells displaying clear, parallel stress fiber bundles were categorized as “stress fiber-positive,” while those with F-actin organized along the cortex of the cells were labeled “cortical.” The percentage of each cell type was calculated relative to the total cell count.

The ratiometric analysis of G- to F-actin (G/F-actin) in cells was examined by tracing cell boundaries in maximum intensity projection confocal images using FIJI ImageJ [14, 26]. The mean fluorescent staining intensity for Vitamin D Binding Protein (G-actin) was divided by the mean fluorescent staining intensity for Phalloidin (F-actin). To account for variances due to staining/imaging G/F-actin for each condition was normalized to the average of the controls for each set of experiments. The normalized data was then pooled.

### KI67 Staining and Quantification

For visualization of KI67, PFA-fixed cells were permeabilized in permeabilization/blocking buffer and then incubated overnight in buffer containing primary antibody against KI67 (1:100; Rabbit polyclonal IgG; ab15580; Abcam). Cells were subsequently washed six times in PBS, and then incubated in secondary antibody solution containing anti-rabbit CF488 (1:100; Biotium), Rhodamine-phalloidin (1:50), and Hoechst (1:500). After a 1 hour incubation at room temperature, cells were washed six times in PBS and then mounted using Everbrite mounting medium. Cells were imaged on a Zeiss LSM880 confocal microscope using a 40x objective.

The proportion of nuclear KI67 was quantified as previously described with slight modifications [14]. Briefly, KI67 fluorescence in the nucleus and cytoplasm was measured by outlining cell and nuclear boundaries on maximum intensity projections of z-stack images using FIJI Image J. Rhodamine phalloidin staining for F-actin at the cell borders served as an indicator of cell boundaries, while Hoechst staining for nuclei served to indicate nuclear boundaries. The raw integrated densities of KI67 signal from the nucleus and area were obtained for each cell and used to determine mean fluorescent cytoplasmic and nuclear intensity. The proportion of KI67 in the nucleus was calculated by ratiometric analysis of nuclear-to-cytoplasmic signal. Due to variances in staining between experiments, the nuclear KI67 for each cell was normalized to the average nuclear KI67 proportion at 36°C. The data obtained from three sets were then pooled.

### Tissue Immunostaining for cartilage matrix molecules

After 20 days in culture, tissues were fixed in 4% PFA at 4°C. Following overnight fixation, tissues were washed with PBS, and placed in 30% sucrose and incubated at 4°C. The next day the tissues were snap-frozen in OCT and cryosectioned to 10-micron sections. Sections were incubated overnight at 4°C with primary antibody against COL2 (1:100; Rabbit; AB761; Millipore, Bedford, MA, USA) and ACAN (1:500; Mouse monoclonal antibody; AHP0022; Invitrogen, Carlsbad, CA, USA). Tissues were then washed in PBS and subsequently incubated in secondary antibody solution consisting of anti-rabbit CF568 (1:200; #20103; Biotium), anti-mouse CF647 (1:200; #20458; Biotium), and Hoechst (1:500). Cells were imaged on a Zeiss LSM880 confocal microscope using a 10x objective.

### Gene expression analysis

RNA from cells and tissues were isolated using TRIzol (Sigma-Aldrich) as previously described [14, 27]. Tissues were homogenized by manually grinding using a Pellet Pestle. Next, chloroform was added into TRIzol solution to allow for phase separation. An RNA Clean Concentrator Kit (Zymo, Irvine, CA, USA) was then used to selectively recover RNA. To reverse transcribe RNA to cDNA a UltraScript 2.0 cDNA Synthesis kit (PCR Biosystems, Wayne, PA) was used. PCR was performed using qPCRBIO Sygreen Blue Mix (PCR Biosystems) according to manufacturers directions and using 20ng of cDNA per reaction. The primers used in this study were previously validated and listed in our previous studies [14, 22]. The PCR reactions were carried out on a Cielo3 real-time PCR machine (Azure; Houston, TX, USA). A melt curve analysis was performed to validate gene produce specificity. Relative mRNA levels were quantified using the deltaCt Pfaffl method using 18S as the reference gene [28].

### Statistical analysis

Experiments were repeated at least three times and conducted on separate occasions from separate animals. Graphpad Prism 9 (San Diego, CA, USA) software was used for statistical analysis of pooled data from experiments. In statistical evaluation, outliers were first determined using the ROUT method with a maximum false discovery rate set at 1% [29]. Differences between groups of data were determined using an analysis of variance (ANOVA) in conjunction with a Dunnett’s post-hoc test.

## Results

### Temperature evaluation of storage environments

To determine the average storage temperatures and temperature fluctuations that occur over time, we evaluated the temperatures within the cell culture incubator, ambient (laboratory) environment, and in the refrigerator. We determined that the incubator, ambient environment and refrigerator temperatures to be 36.03+/-0.06°C (average+/-St.err), 19.42+/-0.06°C+/-, and 7.81+/-0.09°C respectively (Figure S1A). We found only slight variations in temperature ranges for the incubator and ambient environments, but more variation in temperatures in the refrigerator over a 24-hour period (Figure S1B). On average, the temperature ranges spanned 0.85°C, 0.93°C and 2.83°C for the incubator, ambient environment, and refrigerator, respectively. Based on these findings, for the remainder of our study, we referred to the incubator, ambient, and refrigerator conditions as 36, 19, and 8°C, respectively.

### Adhered P2 cells stored under hypothermic temperatures decrease in size and increase in circularity

As a first means to investigate the effect of hypothermic temperatures on passaged chondrocytes, we seeded P2 chondrocytes in 2D. After 24 hours in the 36°C cell culture incubator, cells were either maintained at 36°C, or placed at 19 or 8°C. After 2 hours of hypothermic storage, we observed morphological changes in the cells, which appeared smaller and less spread (Figure 1A). These alterations are more pronounced by 24 hours. We quantified cell area and circularity and found that culturing cells under hypothermic conditions for 24 hours at 19 or 8°C resulted in significantly smaller (Figure 1A, B) and more circular (Figure 1A, C) cells compared to cells cultured at 36 °C.

**Figure 1.**
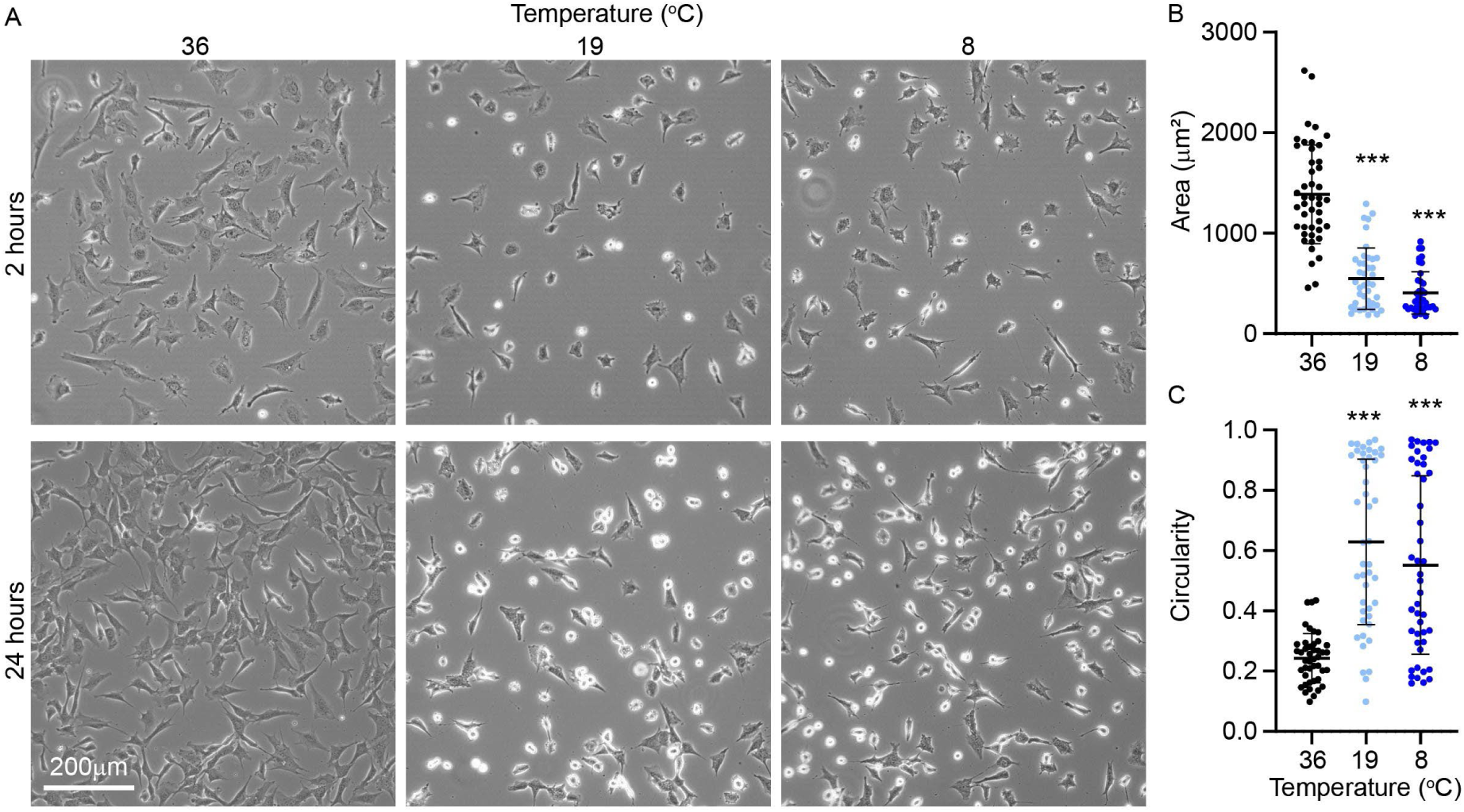
Hypothermia alters morphology of passaged chondrocytes seeded in 2D. (A) Light microscopy images of P2 cells at 36, 19, and 8°C for 2 and 24 hours. Dot plot of image quantification for (B) cell area and (C) circularity of P2 cells. Each dot represents one cell, with data pooled three separate experiments ***, p < 0.001.

### Adhered P2 cells remained viable but do not proliferated after 3 days in hypothermic

Previous studies have demonstrated that hypothermic storage of native cartilage eventually led to chondrocytes death which coincides with cell rounding [18]. Therefore, we investigated the effect of hypothermic temperatures on the viability of P2 chondrocytes in 2D culture. We found that despite the alterations in morphology where hypothermic temperatures caused cell rounding, cells cultured at 19 and 8°C remained highly viable (Figure 2A). After 3 days of hypothermic storage, cell viability is similar to controls as viability was 99.0, 98.6, and 97.4% for cells stored at 36, 19, 8°C for three days, respectively. We observed, however, that culturing under hypothermic temperatures led to less dense cultures. Cellular density after three days is 1.13 x10^5^, 0.50 x10^5^ and 0.33x10^5^ cells/cm^2^ for 36, 19 and 8°C, respectively (Figure 2B).

**Figure 2.**
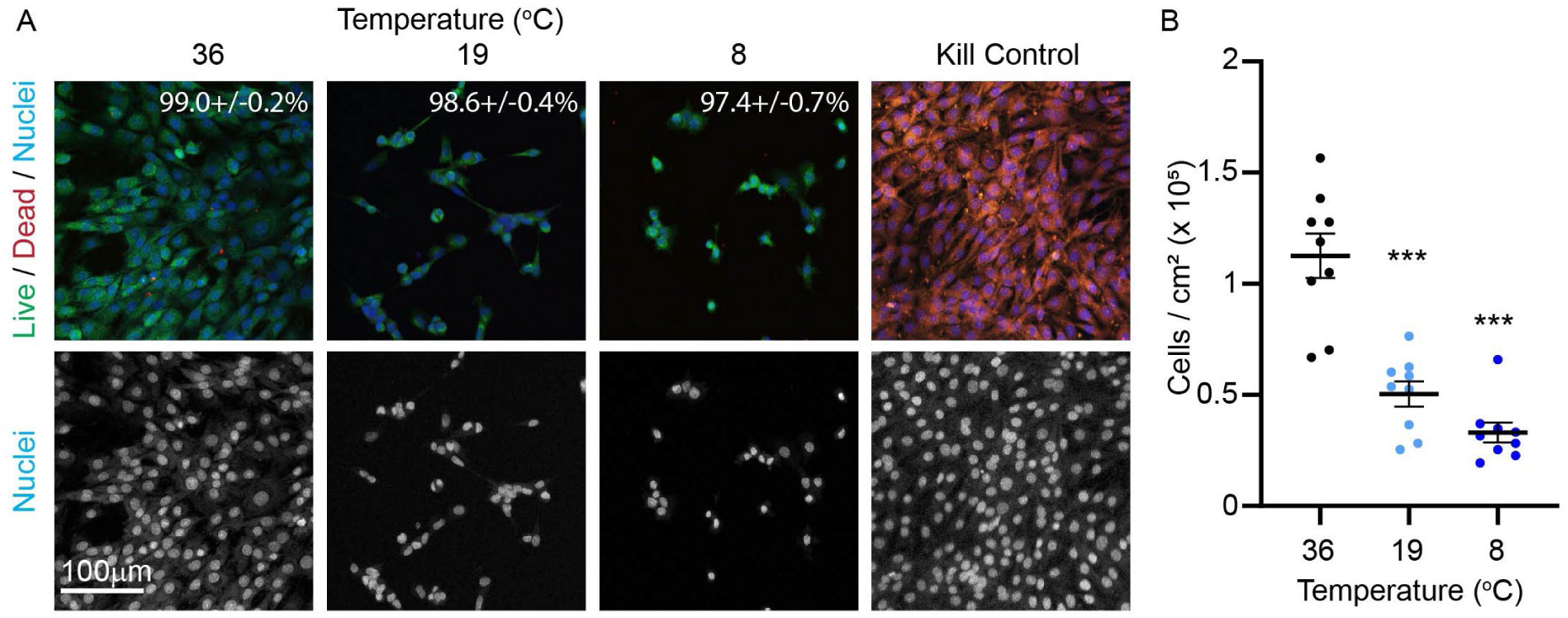
Effect of hypothermic temperatures on P2 cell viability and density in 2D monolayer culture after 3 days of culture. (A) P2 cells were stained for ViaFluor-CF488 (green; live cells), Live-or-dye-CF568 (red; dead cells), and Hoechst (blue; nuclei of all cells). Percent viability was calculated and displayed in top right corner of top panel. (B) Cell density was calculated at various temperatures. ***, p < 0.001 as compared to 36°C

As another means to assess the viability of cells under hypothermic conditions, we evaluated the ability of cells to recover from hypothermia. After 1 day in hypothermia, we allowed the cells to recover at 36°C for 2 days. We found that cells recovered from their rounded state (Figure S2). After 2 days of recovery, cells re-spread (Figure S2A, B). The cell area increased and is similar to that of cells originally maintained at 36°C. The ability to proliferate also recovers as there is a notable increase in cellular density at 48 hours.

### Hypothermic temperatures reduce F-actin stress fibers in adhered P2 cells

Thus far our findings show that cells are viable and round. Round cells are characteristic of primary differentiated chondrocytes. F-actin organization is a determinant of chondrocyte morphology [5, 7, 8]. Previous studies have shown that the storage of non-chondrogenic cells (i.e., foreskin fibroblasts) at hypothermic temperatures causes F-actin depolymerization [30]. Since F-actin is also a key regulator of chondrocyte phenotype and health, we investigated the effect of temperature on F-actin organization and the actin polymerization status. We found that exposure of P2 chondrocytes to either 19°C or 8°C temperatures reduces the proportion of cells that display stress fibers (Figure 3A, B). Instead, a large proportion of cells at 19 and 8°C display cortical F-actin. In addition, by staining for G- and F-actin, we observed an overall increase in staining for both G-/F-actin in cells (Figure 3A, C).

**Figure 3.**
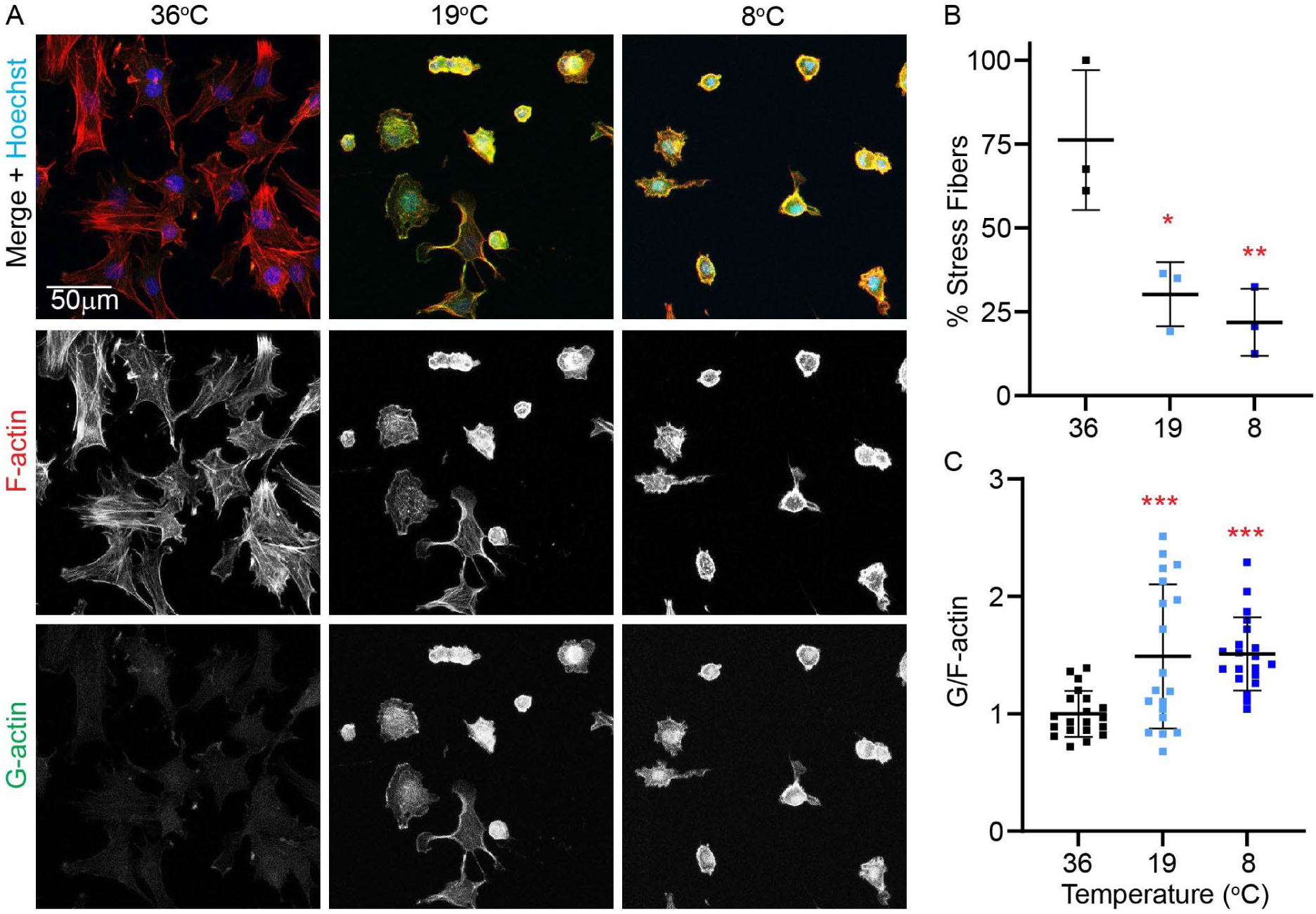
Effect of hypothermic temperatures on actin polymerization. (A) Confocal images of P2 cells stained for vitamin D-binding protein-CF488 (G-actin; green) and Rhodamine-phalloidin (F-actin; red). Cell were counterstained using Hoechst (Nuclei; blue). Images were quantified for (B) proportion of cells that displayed stress fibers, and (C) ratiometric G/F-actin fluorescent intensity. Each dot in (B) represents the proportion of cells with stress fibers from one set of experiments with a total of 3 separate sets of data. Each dot in (C) represents one cell, with data pooled three separate experiments. *, p < 0.05; **, p < 0.01 ***, p < 0.001.

### Hypothermic temperatures reduce the mRNA levels of specific matrix molecules in adhered P2 cells

Storage of passaged cells under hypothermic conditions resulted in viable cells that are round, display cortical F-actin organization, and have higher G/F-actin levels, which resemble the primary chondrocyte phenotype [5, 7, 8]. To further investigate whether storage under hypothermic conditions primes redifferentiation toward a primary-like phenotype, we analyzed gene expression following a 24-hour exposure to hypothermic conditions. We found that hypothermia does not alter the mRNA levels of cartilage matrix protein chondroadherin (*CHAD*) or chondrogenic transcription factor SRY-Box Transcription Factor-9 (*SOX9*) (Figure 4A). However, culture in either 19°C or 8°C reduces cartilage matrix mRNA levels for *ACAN* and *COL2* (Figure 4A). Culture at 8°C also reduces the mRNA levels of fibroblast matrix molecule *COL1* (Figure 4B), but does not alter the mRNA levels of fibroblastic matrix molecule *TNC*. Hypothermic temperatures do not alter the mRNA levels of the fibroblastic isoform of actin, αSMA, or the fibroblast actin stress fiber cross-linker, *TAGLN*.

**Figure 4.**
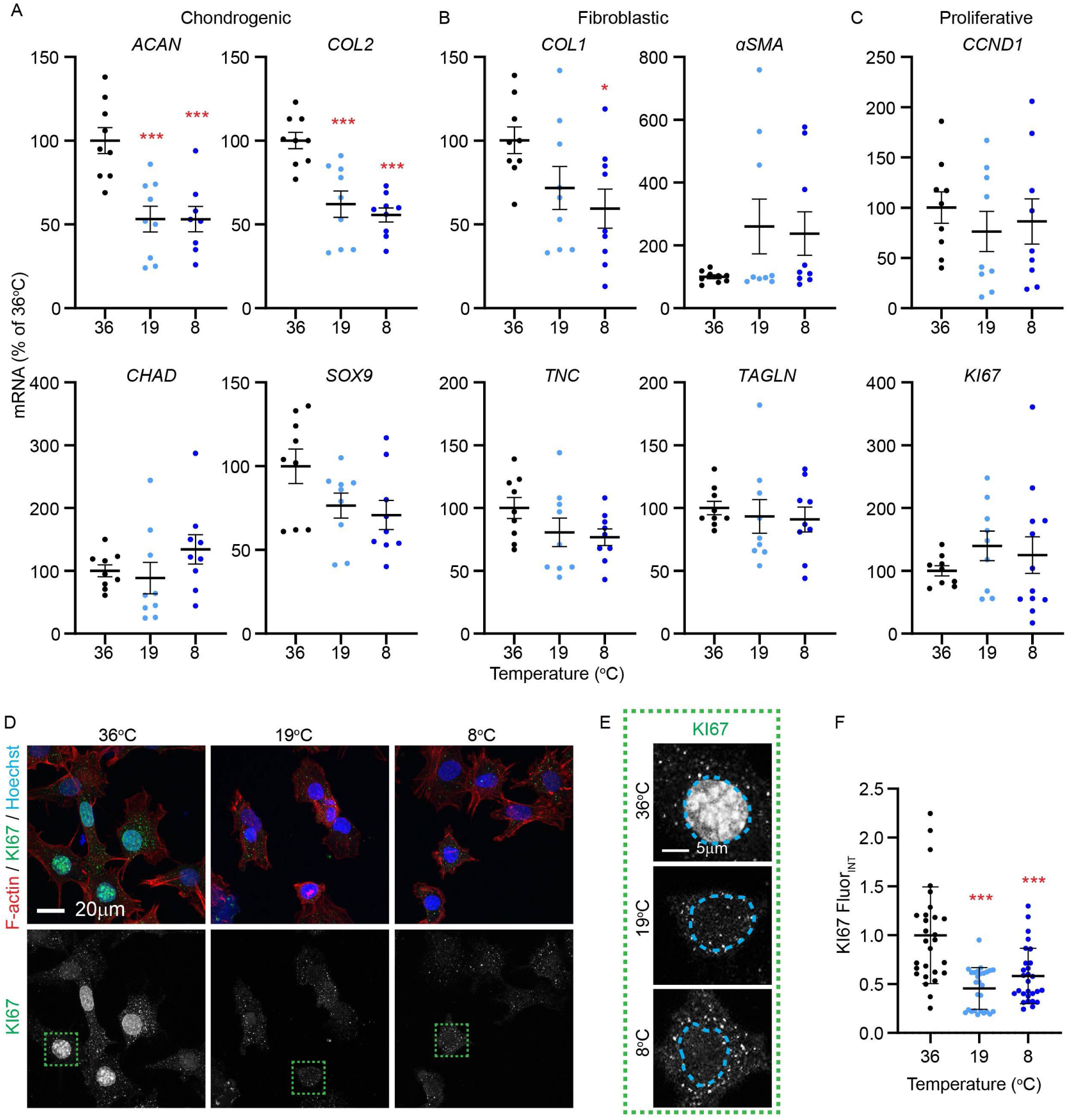
Effect of hypothermic storage on mRNA levels and KI67 nuclear staining of P2 cells in monolayer culture after 1 day of culture. Relative RT-PCR data dot-plots showing mRNA levels for (A) chondrogenic, (B) fibroblastic, and (C) proliferative molecules. (D) KI67 immunostaining (green) of cells. Top panel shows merged image of co-staining with Rhodamine-phalloidin (F-actin) and Hoechst (nuclei; blue). Bottom panel shows separated green channel. (E) Grey scale images taken at higher mag of nuclei outlined in (D). (F) Dot-plot of normalized nuclear KI67 fluorescent intensity proportion. Each dot represents nuclear intensity for one cell, with data pooled three separate experiments. *, p < 0.05; ***, p < 0.001 as compared to 36°C control.

### Hypothermic storage reduces nuclear localization of Ki67

Since hypothermic storage of cells led to a lower cell density, we also investigated the mRNA levels of critical chondrocyte proliferative molecules, cyclin D1 (CCND1) and the proliferation marker Ki-67 (KI-*67*). We found that hypothermic conditions do not affect the mRNA levels of either *CCND1* or *KI67* (Figure 4C). However, since actin reorganization/depolymerization can also alter the proportion of nuclear KI67 [14], we next stained cells cultured in hypothermic conditions for KI67. We found that compared to culturing cells at 36°C, culture in either 19 or 8°C reduces the proportion of nuclear KI67 (Figure 4D-F).

### Effect of temperature on morphology and viability of P2 cells in suspension

Since cells for ACI may be shipped from the laboratory to the clinic in suspension [31], we next investigated the response of P2 cells in suspension to hypothermic temperatures. Light microscopy revealed that cells in suspension were rounded and small at all temperatures as compared to attached cells (Figure 5A). We found that after 24 hours, P2 cells in suspension maintained at 36°C formed aggregates, whereas those stored at 19 or 8°C did not. Cell death within aggregates, as well as in single cells that remained in suspension, is visible. By performing viability analysis, we found that cells maintained at 36°C for 1 day are less viable, both in aggregates as well as single suspended cells, than those stored at 19 or 8°C for 1 day. After 3 days of culture, P2 cells in suspension, maintained at 36°C, showed a further reduction in cell viability. The P2 cells in suspension are more viable when maintained at 19 or 8°C for 3 days. The viability of cells at 19 and 8°C after 3 days are similar to the respective viabilities at 1 day.

**Figure 5.**
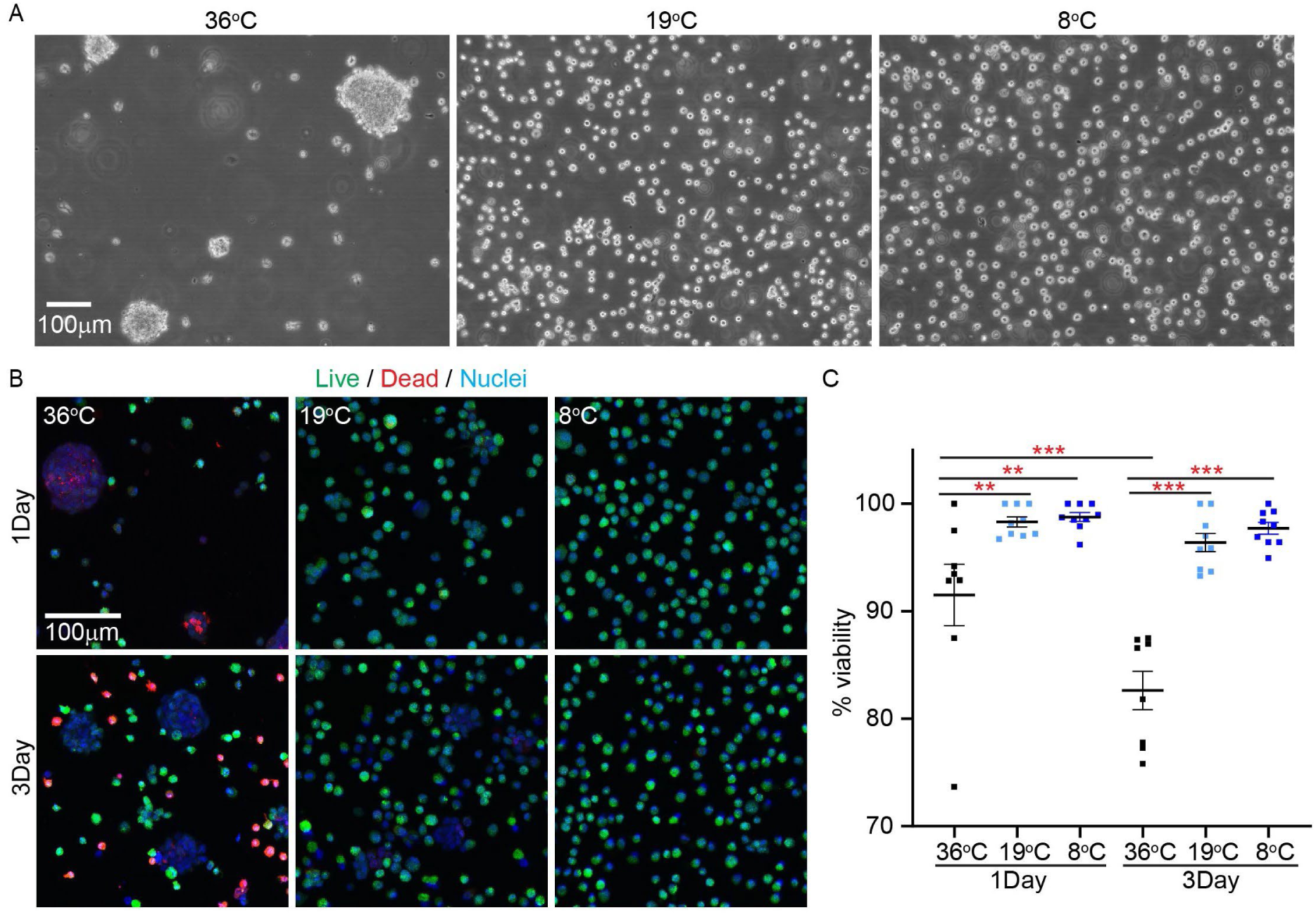
The effect of hypothermic temperatures on P2 cell aggregation and death in suspension culture. (A) Light microscopy images of cells in suspension culture. (B) Cells in suspension stained for ViaFluor-CF488 (green; live cells), Live-or-dye-CF568 (red; dead cells), and Hoechst (blue; nuclei of all cells). (C) Dot-plot showing calculated percent viability at various temperatures. **, p < 0.01; ***, p < 0.001 as compared to 36°C.

### Effect of temperature on mRNA levels of P2 cells in suspension

We next evaluated the effect of temperature on P2 cell mRNA levels in suspension. As compared to adhered P2 cells on polystyrene at 36°C, P2 cells in suspension at 36°C elevate *CHAD* mRNA levels (Figure 6A). However, there are no significant differences caused by culturing cells in suspension at 36^ο^C for *ACAN*, *COL2*, and *SOX9*. However, while in suspension, culture of P2 cells at 8°C led to a specific upregulation of *ACAN* and *COL2* mRNA levels as compared to P2 cells in suspension at 36°C. There are no differences in the expression of *CHAD* and *SOX9* by culture of P2 cells in suspension at 8°C versus 36°C. We found that culture P2 cells in suspension at 19°C results in similar mRNA levels of *ACAN*, *COL2*, *CHAD*, and *SOX9* as compared to P2 cells in suspension at 36°C.

**Figure 6.**
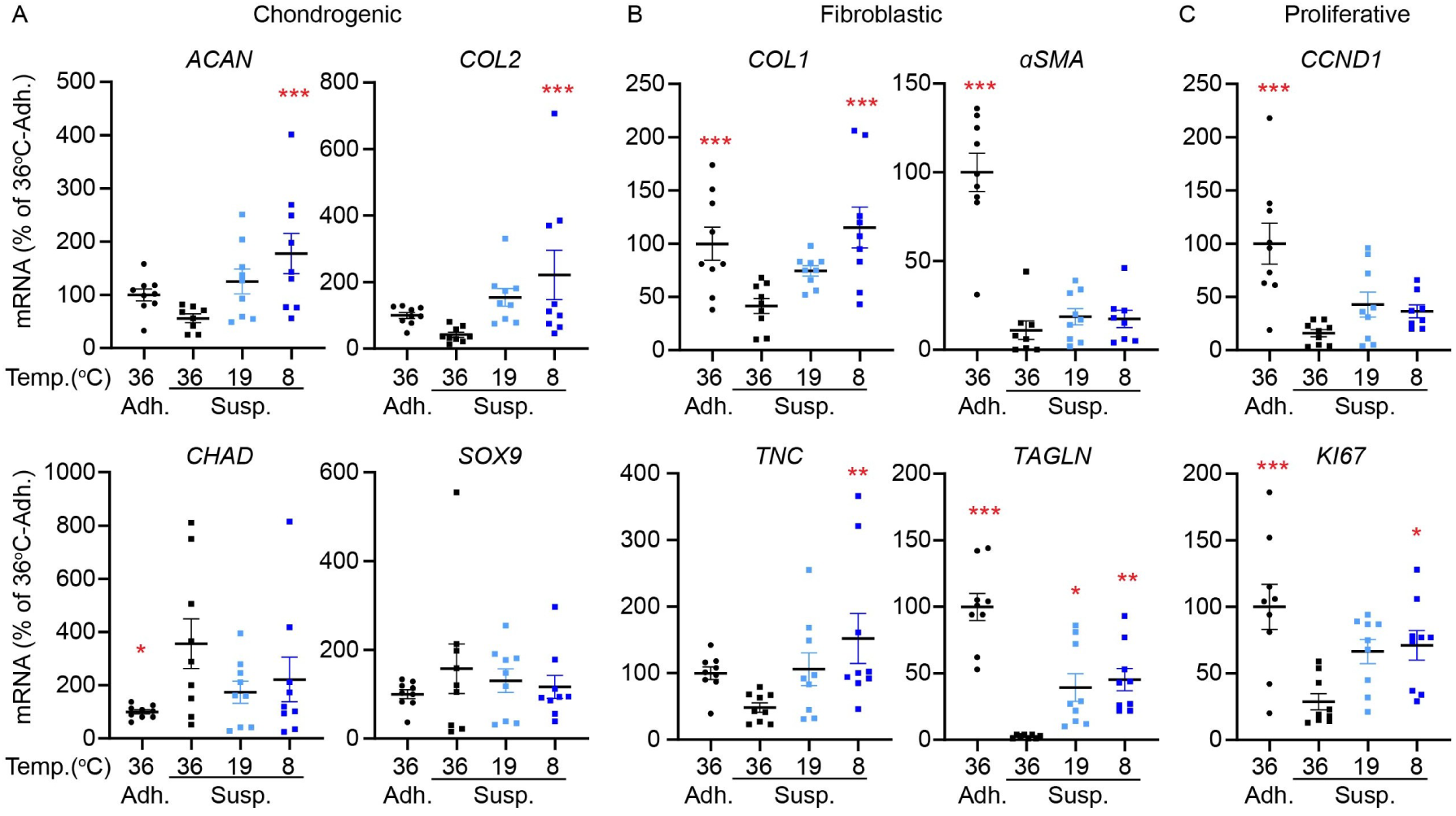
Relative real-time RT-PCR of P2 cells cultured in suspension at hypothermic temperatures for 1 day. (A) Chondrogenic, (B) fibroblastic, and (C) proliferative molecule mRNA levels of P2 cells in suspension at hypothermic temperatures as compared to P2 cells in suspension. As a reference point, mRNA levels were normalized to control cells that were adhered in monolayer culture and stored at 36°C. **, p < 0.01; ***, p < 0.001 as compared to cells in suspension stored at 36°C.

With regard to the expression of fibroblastic matrix and contractile mRNA levels, compared to adhered P2 cells at 36°C, the culture of cells in suspension at 36°C decreases *COL1*, *αSMA*, and *TAGLN* (Figure 6B). There was no significant difference in *TNC* mRNA levels between adhered cells and suspension cells at 36°C. As compared to P2 cells in suspension at 36°C, the mRNA levels for *COL1*, *TNC*, and *TAGLN* are elevated by culture of P2 cells in suspension at 8°C. There is a similar upregulation in the mRNA levels for *TAGLN* following the culture of P2 cells in suspension at 19°C. Unlike the culture at 8°C, P2 cells in suspension at 19°C have similar mRNA levels for *COL1* and *TNC* as compared to P2 cells in suspension at 36°C. Neither 19 or 8°C, impacted the *αSMA* mRNA levels as they are similar to P2 cells in suspension at 36°C.

Finally, with regard to expression of proliferation molecules, culture of P2 cells in suspension at 36°C led to a reduction in *CCND1* and *KI67* mRNA levels as compared to adhered cells at 36°C (Figure 6C). Storage of P2 cells in suspension at 8°C had higher *KI67* mRNA levels than in P2 cells in suspension at 36°C. Culture of P2 cells in suspension at 8°C, however, did not affect *CCND1* mRNA levels. Culture of P2 cells in suspension at 19^ο^C did not affect *CCND1* or *KI67* mRNA levels.

### Effect of hypothermic temperatures on P2 chondrocyte redifferentiation and matrix formation in 3D cultures

Culture of P2 cells in suspension at hypothermic conditions promotes viability, reduces cell aggregation, and enhances the mRNA levels of specific matrix molecules including *ACAN* and *COL2*. This led us to speculate that maintaining P2 cells in suspension at hypothermic temperatures may benefit the production of cartilage matrix by P2 cells. To determine the effect of acute hypothermia on the longer-term ability of P2 cells that were cultured in suspension to form a cartilaginous matrix, we maintained P2 cells in suspension for a period of 1 day prior to seeding cells in 3D adherent agarose mold cultures for the generation of scaffold-free cartilage tissue [22, 23]. After 1 day in 3D culture, we found that P2 cells from suspension culture at 36°C form dense aggregates (Figure 7A), which we do not see in P2 cells in 3D from adherent cultures at 36°C. We found that P2 cells from suspension at 19 or 8°C do not form aggregates.

**Figure 7.**
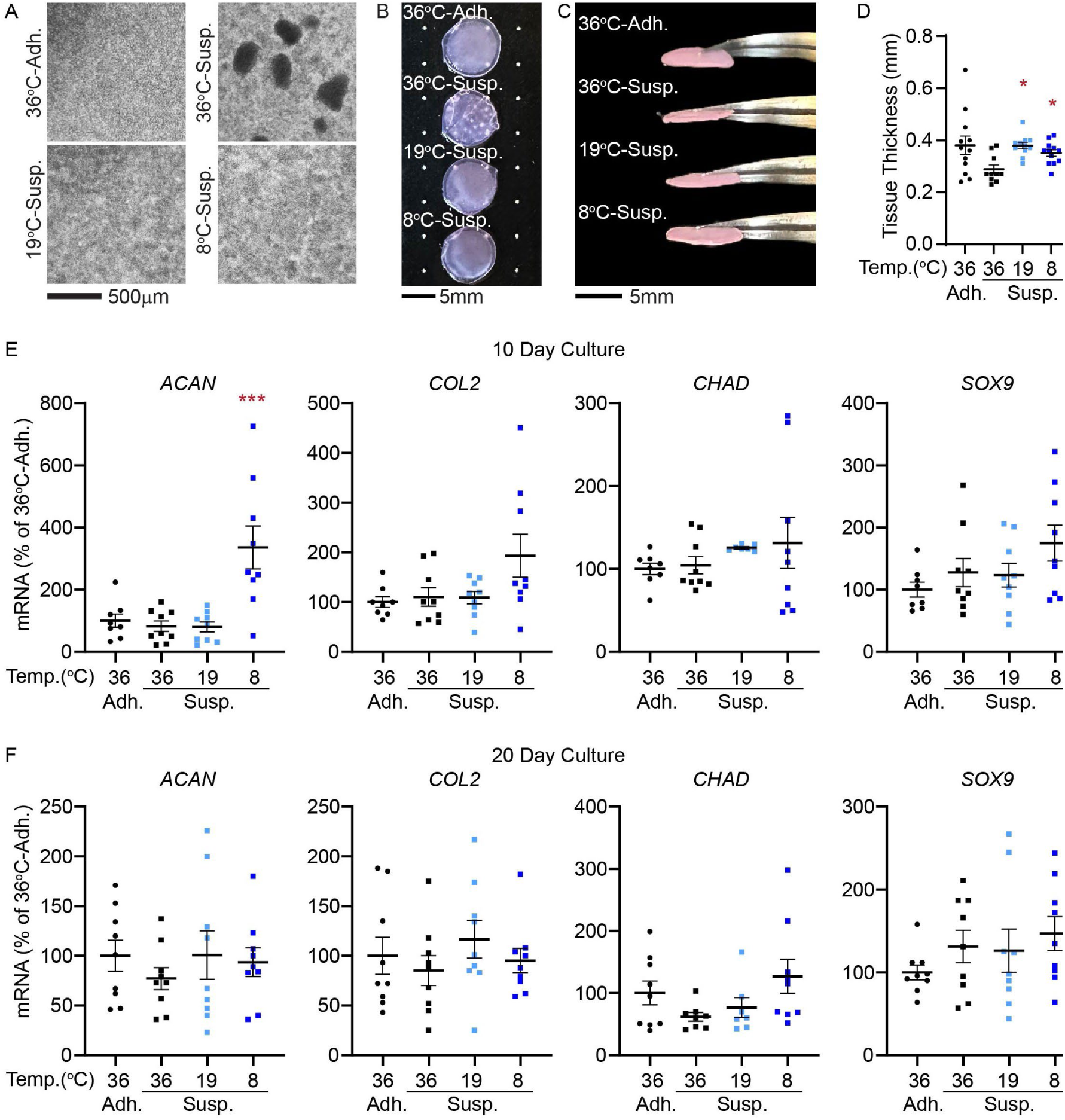
P2 cell ability to redifferentiate in 3D adAM cultures following storage in monolayer at 36°C, or in suspension at 36, 19, or 8°C. (A) Light microscopy images of P2 cells after 1 day in adAM cultures. (B) Top view images of tissues on situated on dotted graphing paper after 20 days of adAM culture. (C) Side view images of tissues being held by forceps after 20 days of adAM culture. (D) Measured tissue thickness of constructs after 20 days of adAM culture. Relative RT-PCR mRNA levels for chondrogenic molecules after (E) 10 and (F) 20 days of adAM culture. As a reference point, mRNA levels were normalized to control cells in adAM culture that were derived from adhered monolayer culture at 36°C. *, p < 0.05; **, p < 0.01; ***, p < 0.001 as compared to cells in suspension stored at 36°C.

**Figure 8.**
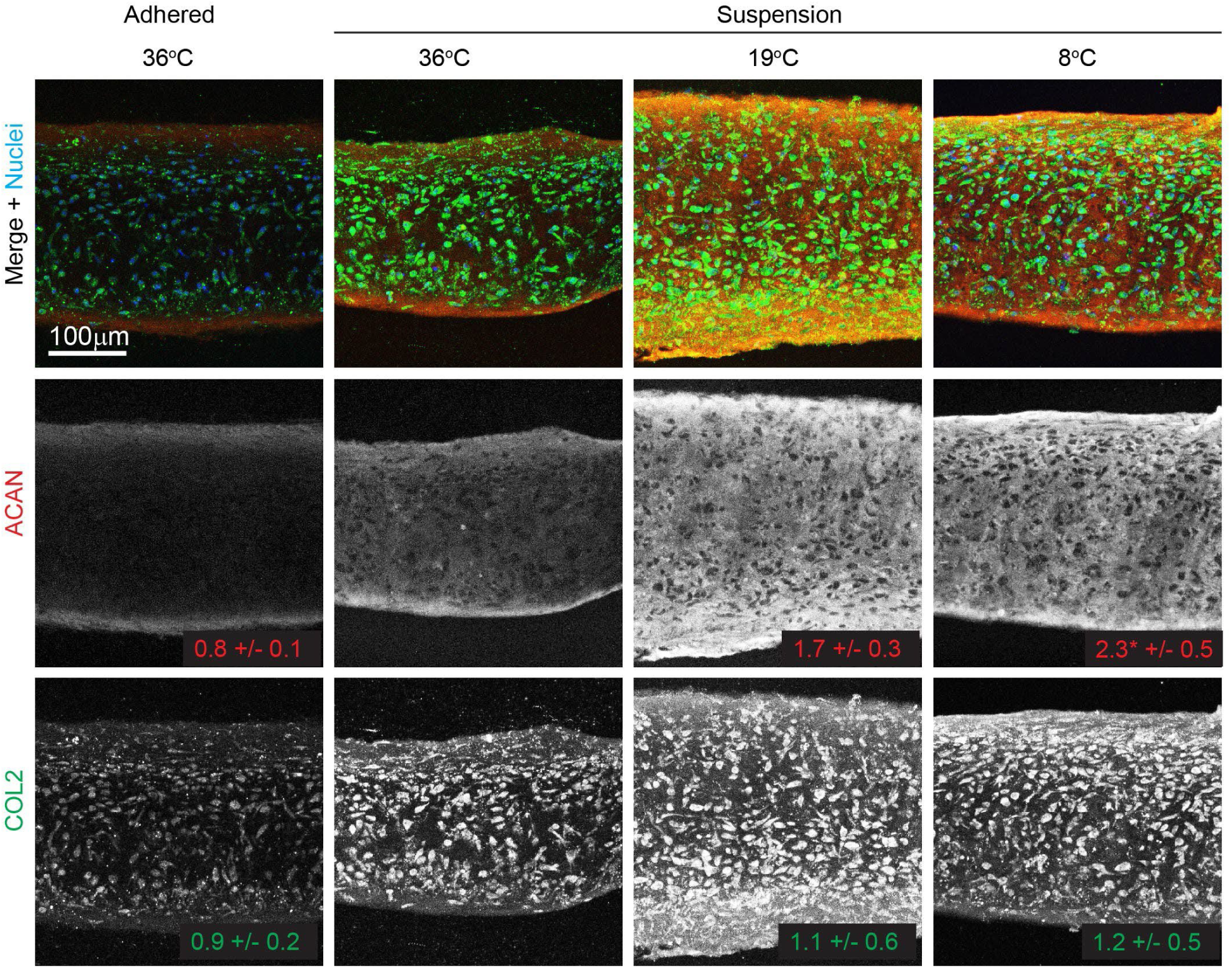
The effect of storage in monolayer at 36°C, or in suspension at 36, 19, or 8°C, on P2 cell ability to produce cartilage matrix in 3D adAM cultures. Sections of 20day adAM tissues were stained for ACAN (red), COL2 (green), and nuclei (blue). Top panels show merged images of all stains, middle shows separated ACAN channel in grey scale, and bottom shows separated COL2 in grey scale. The numbers at the bottom right of ACAN and COL2 images represents the relative average staining intensity of each condition for ACAN and COL2 as compared to cells that were stored in suspension at 36°C. A total of six tissues (N = 6) were sectioned and stained from separate experiments. *, p < 0.05 as compared to cells in suspension at 36°C.

P2 cells in 3D culture form matrix, irrespective of their the temperature they were stored in. We found that after 20 days, cells in suspension at 36, 19, and 8°C formed matrix (Figure 7B) that are stable, maintain their disc structure, and can be handled using forceps (Figure 7C). While we found there to be no significant differences (p = 0.08) in tissue thickness between P2 cells in suspension that are maintained at 36°C as compared to control P2 cells from adhered culture (Figure 7D), we did observe a significant difference when P2 cells in suspension are maintained under hypothermic conditions. P2 cells in suspension that were maintained at 19 and 8°C prior to seeding in 3D develop thicker tissues as compared to P2 cells in suspension at 36°C. In addition to differences in tissue thickness, we observed in top view images of tissues (Figure 7B) that acute storage of P2 cells in suspension at 36°C prior to seeding in 3D cultures, results in the formation of dense opaque plaques. These plaques are absent when cells in suspension are stored at 19 or 8°C prior to seeding in 3D culture. The formation of dense plaques is consistent with the cell aggregates formed early in 3D culture by P2 cells in suspension at 36°C (Figure 7A).

### P2 cells from suspension culture at 8°C form cartilaginous tissue with greater staining for ACAN

To evaluate the production of cartilage specific molecules by P2 cells stored in suspension at various temperatures, the tissues formed by cells in 3D culture at 20 days were sectioned and stained for the cartilage matrix molecules, ACAN and COL2. We found that P2 cells stored in suspension at 36, 19, or 8°C all formed tissue that are positive for ACAN and COL2. The storage of P2 cells in suspension at 36°C had similar staining than P2 cells from adhered cultures at 36°C. There is no significant effect of storing P2 cells in suspension at 19°C, as compared to storing P2 cells in suspension at 36°C, on ACAN or COL2 staining. Furthermore, there is no significant effect of storing P2 cells in suspension at 8°C, as compared to storing P2 cells in suspension at 36°C, on COL2 staining. However, by storing P2 cells in suspension at 8°C, there is greater staining for ACAN as compared to storing P2 cells in suspension at 36°C. Storing P2 cells in suspension at 8°C for 1 day prior to 3D culture has a positive effect on the matrix formation by cells.

## Discussion

This study tested the hypothesis that storing passaged cells at hypothermic temperatures would negatively impact cell viability, as well as their capacity to redifferentiate and form matrix. Our findings, however, contradicted our hypothesis. We found that culturing passaged cells under hypothermic temperatures for up to three days did not negatively impact the cellular viability of cells in either monolayer or suspension culture. Rather, we found that the hypothermic storage of passaged cells maintained in suspension culture surprisingly promoted cell viability. In addition, while the culture of P2 cells in 2D monolayer on polystyrene results in a slight reduction in the expression of cartilage matrix molecules, we found that passaged cells stored in suspension culture at hypothermic temperatures exhibit elevated matrix molecule mRNA levels. Furthermore, the tissues produced by passaged cells that were maintained at 9^°C^ produced thicker tissue that stained more intensely for ACAN. These results demonstrate that storing cells under hypothermic temperatures for cell transplantation purposes is not detrimental and can be beneficial for maintaining passaged cell viability and the production of cartilaginous matrix.

The culture of cells under hypothermic temperatures was not detrimental to cell viability in 2D culture, and hypothermic temperatures promoted viability up to 3 days in suspension. This is in contrast to findings on non-chondrogenic mammalian cells (M2, 293T, SaOS2, and mESC) at cooler temperatures (i.e. 4°C) reduces viability [21]. Our findings also contrast with previous studies on chondrocytes in native cartilage [16, 17, 32, 33]. It was shown that incubating biopsied discs of human knee cartilage at 22°C resulted in greater chondrocyte death than when cultured at 37°C [17]. The finding of reduced chondrocyte viability under hypothermic temperatures is also found when osteochondral samples from humeral heads of goats are stored at 4^ο^C [16]. Notably, we only stored cells for up to 3 days at hypothermic temperatures. We chose up to 3 days as our time point for investigation, as cells shipped for transplantation are typically used within this 3-day timeframe [31]. Studies involving the storage of native cartilage in hypothermic temperatures often assess viability after an extensive culture period (i.e., 4 weeks) [16, 17, 32, 33]. It is plausible that hypothermic storage could lead to cell death in passaged chondrocytes if stored for more extended periods; however, this was not assessed in this study. Additionally, the environmental conditions in which the cells exist may also play a role in regulating their viability. Cells within native cartilage may behave differently than cells in monolayer or suspension culture, as they are influenced by reduced interaction with matrix molecules and increased cell-cell interactions.

The *in vitro* cell context, whether stored in a monolayer or suspension, impacts cell viability. While passaged cells maintained in monolayer culture at 36°C were viable, we found that passaged cells maintained in suspension at 36°C were less viable. Cell death in suspension may be caused by the loss of cell attachment in the process of anoikis [34]. The relationship between anoikis and temperature has not been completely elucidated, however, temperature can affect the interaction between integrins and ligands [35]. Therefore, it is possible that temperature modulates the effect of loss of cell attachment by regulating integrin signaling. In addition to anoikis, it is possible that the decrease in cell viability of passaged chondrocytes in suspension at 36°C is due to the formation of cellular aggregates which may prevent the flow of nutrients, or oxygen to cells. Since cooler temperatures prevented aggregate formation, they may be more able to access nutrients and/or oxygen leading to greater viability. However, we also found that cells not in aggregates to be dead. Therefore, aggregate formation may not explain all the death that occurs in suspension cultures at 36°C. The mechanism(s) by which hypothermic temperatures prevent cell death in suspension remain unclear. The influence of temperature on cell viability remains to be interesting as hypothermia can be chondroprotective and prevent induced chondrocyte death after trauma [36]. A further understanding on the temperature regulated mechanisms that mediate cell viability could lead to new methods to prevent chondrocyte death in various scenarios.

Hypothermic temperatures slow the proliferation of passaged cells when cultured in 2D monolayer. This finding is consistent with observations in other cell types, where culture under cooler temperatures has been shown to reduce the growth rate of cells. Reducing the culture temperature of CHO cells from 37 to 33°C reduces growth rate [37]. In addition, cold stressing rats by housing at temperatures between 6-8°C, instead of room temperature (∼25°C), reduced epithelial cell proliferation [38]. By reducing the culture temperature, cells in S phase complete synthesis and arrest in G1 phase [39]. Hypothermic temperatures have been shown to regulate critical regulators of proliferation. In a previous study, it was found that culturing human adipose cells for 1 day in a hypoxic environment at hypothermic temperatures (30°C) reduced *KI67* mRNA levels as compared to culturing under hypoxic conditions alone [40]. However, in passaged chondrocytes, we found that cooler temperatures did not affect the expression of the proliferation molecules, *KI67* or *CCND1*. We did observe that hypothermia affected the KI67 in the nucleus of cells, which is a novel finding. Coinciding with a reduction in nuclear KI67, we found that cells in monolayer at hypothermic temperatures had elevated staining for G/F-actin which suggests that hypothermic temperatures promote actin depolymerization. In a previous study, we found that the depolymerization directly reduces the proportion of nuclear KI67 [14]. Therefore, the regulation of KI67 by hypothermia may be through actin-mediated processes.

Hypothermic storage of passaged cells in 2D repressed matrix molecules (*ACAN, COL2, COL1)* mRNA levels, which also coincided with actin depolymerization. The reduction in *ACAN* is in keeping with findings in primary chondrocytes and osteochondral allografts where hypothermic storage reduces proteoglycan content [19, 41]. Furthermore, albeit at much colder (cryopreservation) temperatures, freezing chondrocytes within a hyaluronan scaffold reduces *COL2* expression [42]. The reduction in expression for *ACAN, COL2,* and *COL1* may also be explained by actin depolymerization. In previous studies, we have found that actin depolymerization reduces the expression of these molecules in passaged chondrocytes [11, 14, 23]. Intriguingly, we did not observe a reduction in matrix molecule expression following hypothermic storage of passaged cells in suspension; instead, we found an increase in matrix molecule mRNA levels at 9°C compared to cells in suspension at 36°C. This speaks to the context-specific effect of hypothermia on matrix molecule expression. Notably, the culture of cells in suspension or in 3D culture results in a reduction of F-actin, which initially leads to a decrease in matrix molecule expression [11, 23]. In support of this, we observe a reduction in COL1 mRNA levels when passaged cells are cultured in suspension at 36°C, as compared to adherent cultures at 36°C. The levels for *COL2*, *ACAN*, and *TNC* also trended toward downregulation in culture suspension; however, these were deemed not statistically significant. We have previously found that in 3D cultures, the levels of chondrogenic matrix molecules can be recovered by culturing in redifferentiation media [23]. Therefore, while actin depolymerization can initially downregulate the expression of matrix molecules, other independent mechanisms can compensate and induce the upregulation of chondrogenic matrix expression. The positive effect of hypothermic storage on enhancing cartilage matrix deposition is also exemplified in pellet cultures where hypothermic storage at 32°C results in greater wet weight of matrix after 21 days. The effect of hypothermic storage on passaged chondrocytes is context-specific and complex.

Intriguingly, the acute hypothermic storage of passaged chondrocytes in suspension at 9°C had lasting favorable effects on matrix production *in vitro*. In a previous study investigating the effect of hypothermic storage on the redifferentiation of passaged chondrocytes, it was shown that culturing passaged chondrocyte pellets at 32°C, compared to 37°C, suppressed redifferentiation [43]. However, in this previous study, cells were redifferentiated for an extended period (i.e., up to 42 days in culture). In our current study, passaged cells were maintained in suspension at hypothermic temperatures for only 1 day before seeding in 3D culture. Following seeding, the 3D cultures were maintained at 36°C. Another difference is that the hypothermic temperatures used in our current study were also lower, at 19°C and 8°C, compared to the 32°C previously used [43]. We also observed more favorable effects of hypothermic storage when passaged cells in suspension were stored at the 8°C temperature. While tissue thickness was greater for cells that were stored in suspension at 19°C as compared to 36°C, the chondrogenic mRNA levels and staining for cartilage matrix molecules are similar between 19°C and 36°C. However, storage of passaged cells in suspension at 8°C, resulted in thicker tissue, an enhancement of *ACAN* mRNA levels at 10 days of redifferentiation, and greater staining for ACAN after 20 days of redifferentiation as compared to passaged stored in suspension at 36°C. The mechanisms by which colder temperatures may induce favorable matrix deposition is unclear. However, hypothermia has been shown to regulate cold-shock proteins that can affect matrix health. For instance, hypothermia has been shown to upregulate the expression of RNA-binding motif protein-3 (RBM3) in osteoblasts [44]. While there is a limited understanding on the regulation of RBM3 in chondrocytes, it is expressed in almost all tissues. In osteoblasts, RBM3 promotes osteoblastic differentiation through activation of ERK activity [44]. ERK is a known regulator of ACAN expression in chondrocytes [45]. It would be of interest to investigate the regulation of ACAN expression by RBM and ERK, and the regulation differences at 19 and 8°C.

In addition to the differences in matrix production, hypothermic temperatures repressed aggregate formation in suspended passaged cells. The inhibition of aggregation by hypothermic storage is in line with previous findings where resuspending hamster fibroblast BHK21 cells following trypsinization form aggregates at 37°C but not at 2°C[46]. At low temperatures cell-cell adhesion is impaired due to several factors such as reduced protein-protein interactions, adhesion molecules (i.e. cadherins) are less stable/functional, and low temperatures reduce membrane fluidity disrupting integrin-ligand interactions [47–49]. The aggregation of passaged cells in the present study appears to result in the deposition of dense opaque matrix plaques. It is unclear whether these dense plaques cause regional variations in micromechanical tissue characteristics. However, these plaques are absent in tissues formed by cells stored under hypothermic conditions which have more uniform matrix production.

Acute, short-term hypothermic storage of passaged cells, particularly in suspension culture, conferred several benefits that may be favorable for transplantation. Indeed, there are limitations of our study that must be considered. One confounding variable that must be considered was that the 8°C temperature, which had the most favorable effects, was maintained by storing cells in a refrigerator where temperatures had greater fluctuations than storage at ambient, room temperature (19°C) and in a culture incubator (36°C). There is a limited understanding on the effect of temperature fluctuation on cellular health. A second confounding variable is that hypothermic temperatures did not control for CO2 or humidity, as would be controlled in a cell culture incubator. The role that such factors play in the temperature response is yet unclear. Finally, transporting cells from laboratory o the clinic for cell-based transplantation would expose the cells to other stimuli such as mechanical stimulation from package handling and/or transport. The interaction between mechanical stimuli and hypothermic storage of passaged chondrocytes were not considered in this study.

Nevertheless, hypothermic storage of passaged chondrocytes could be favorable for tissue formation in cell transplantation procedures. Passaged chondrocytes stored in suspension culture at 9°C produce thicker tissue that is homogenous and contains higher expression of ACAN. These promising findings warrant further investigation to determine whether the beneficial effects of hypothermic storage on tissue formation *in vitro* are retained in an *in vivo* model.

## Acknowledgments

The research reported in this project was supported by the Delaware Center for Musculoskeletal Research from the National Institutes of Health’s National Institute of General Medical Sciences under grant – NIGMS (P20 GM139760). This publication was made possible by the Delaware INBRE program, supported by a grant from the National Institute of General Medical Sciences – NIGMS (P20 GM103446) from the National Institutes of Health and the State of Delaware.

---

**Figure S1.**
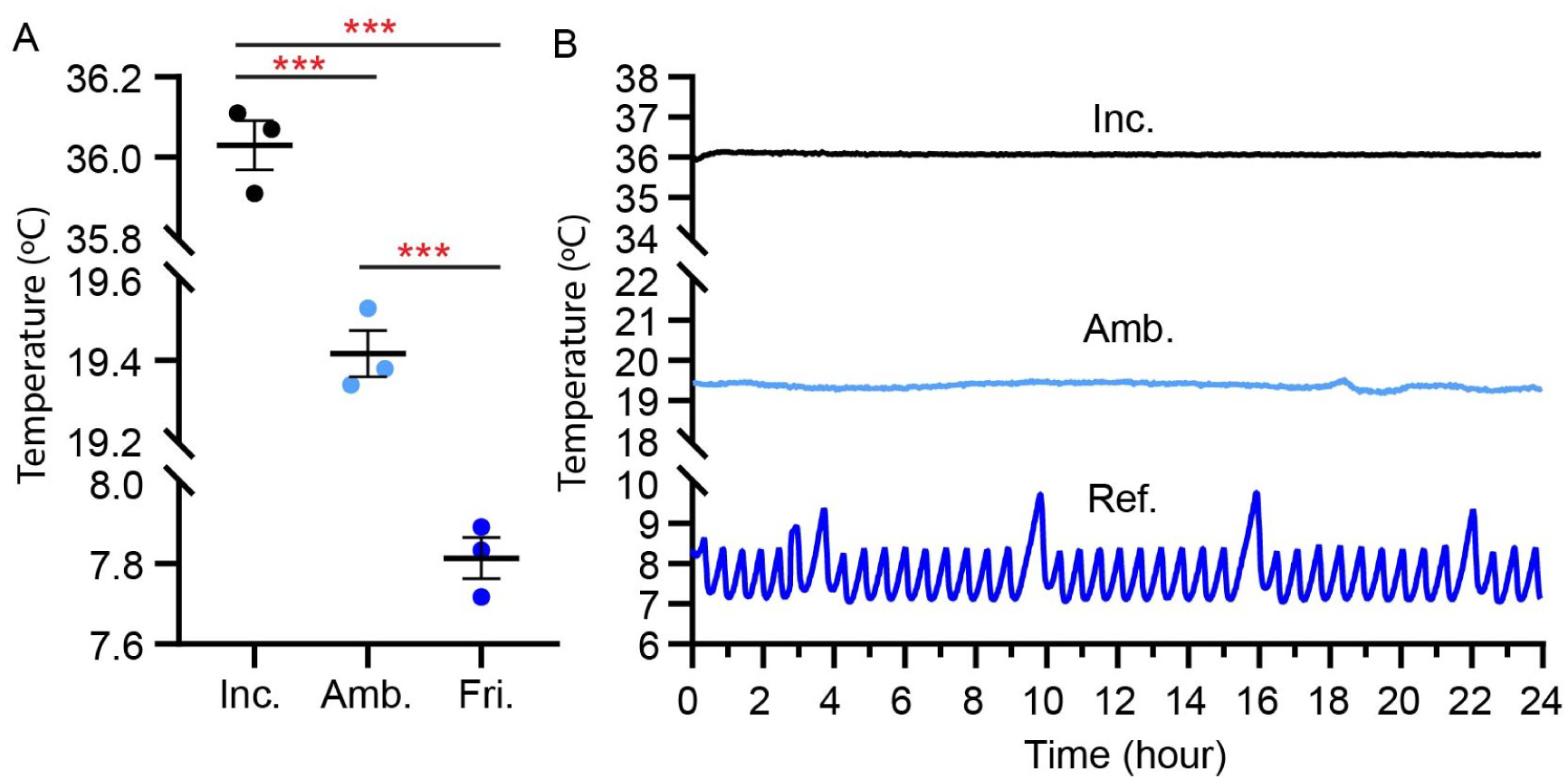
Temperature measurements of CO2 incubator (Inc.), ambient room (Amb.) and refrigerator (Ref.) environments. (A) Average temperature measured over the span of 24 hours. Each point represents the mean temperature measured on three separate occasions. (B) Representative line graph of temperature fluctuations over 24 hours. ***, p < 0.001 as compared to Inc.

**Figure S2.**
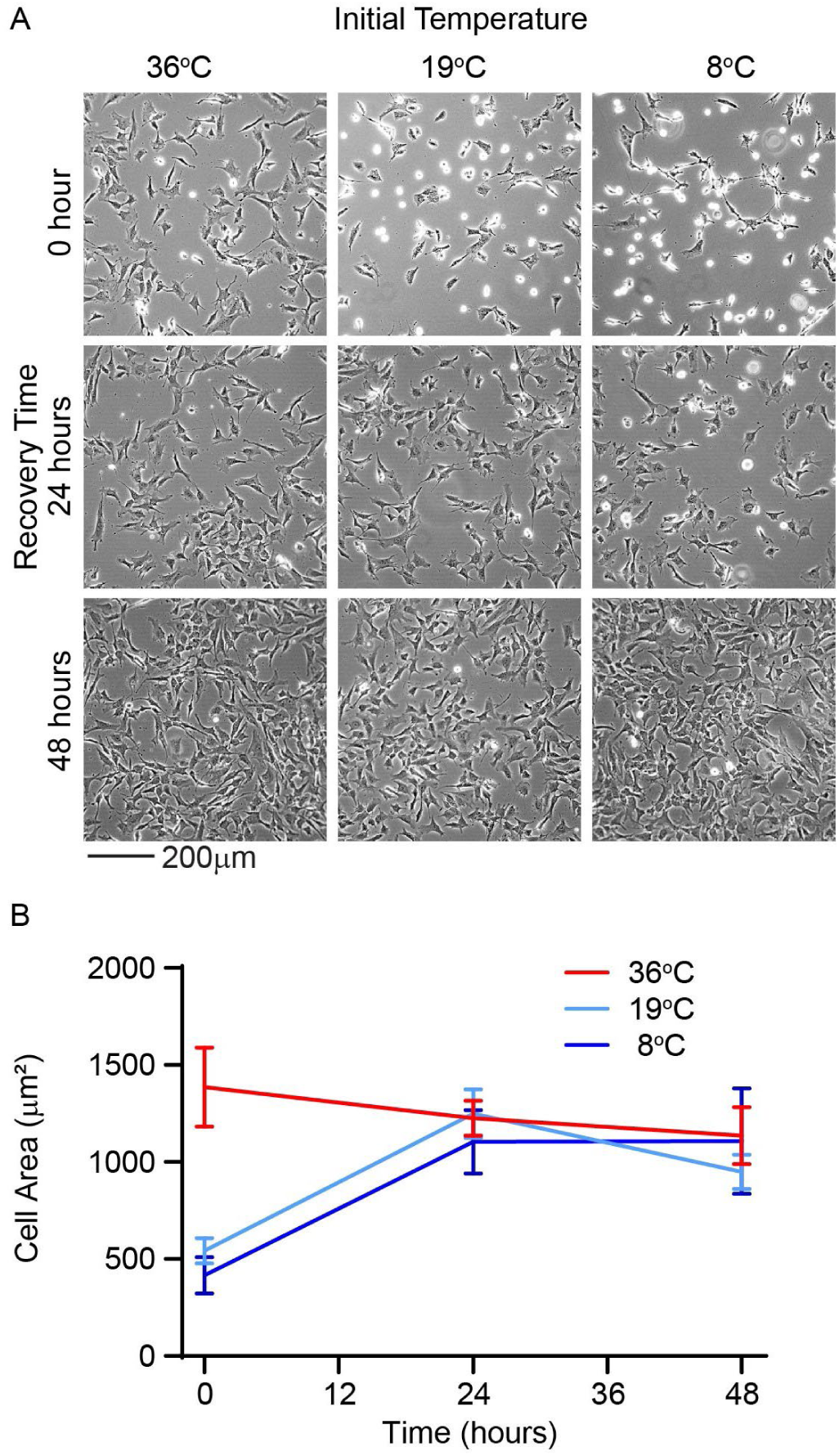
Light microscopy of P2 cells in monolayer culture recovering at 36°C. (A) Images of recovering cells from 36, 19, and 8°C temperatures at 0, 24, and 48 hours post-recovery. (B) Line graph demonstrating that cell area from hypothermic temperatures is recovered by 24 hours.

**Table S1.** Media conditions used in redifferentiation study.

| Culture Day | Media Type (%) |  | Supplement |
| --- | --- | --- | --- |
|  | Complete | Redifferentiation |  |
| 0 | 100 | 0 | Y27632 |
| 1 |  |  |  |
| 2 | 50 | 50 | Y27632 |
| 3 |  |  |  |
| 4 | 0 | 100 | Y27632 |
| 5 |  |  |  |
| 6 | 0 | 100 | Y27632; TGFβ3 |
| 7 |  |  |  |
| 8 | 0 | 100 | Y27632; TGFβ3 |
| 9 |  |  |  |
| 10 | 0 | 100 | TGFβ3 |
| 11 |  |  |  |
| 12 | 0 | 100 | TGFβ3 |
| 13 |  |  |  |
| 14 | 0 | 100 | TGFβ3 |
| 15 |  |  |  |
| 16 | 0 | 100 | TGFβ3 |
| 17 |  |  |  |
| 18 | 0 | 100 | TGFβ3 |
| 19 |  |  |  |
| 20 | 0 | 100 | TGFβ3 |

## References

[1] M. Brittberg, A. Lindahl, A. Nilsson, C. Ohlsson, O. Isaksson, L. Peterson, Treatment of deep cartilage defects in the knee with autologous chondrocyte transplantation, N Engl J Med 331(14) (1994) 889–95.

[2] D.A. Grande, M.I. Pitman, L. Peterson, D. Menche, M. Klein, The repair of experimentally produced defects in rabbit articular cartilage by autologous chondrocyte transplantation, J Orthop Res 7(2) (1989) 208–18.

[3] D.A. Grande, I.J. Singh, J. Pugh, Healing of experimentally produced lesions in articular cartilage following chondrocyte transplantation, Anat Rec 218(2) (1987) 142–8.

[4] J.L. Cook, J.P. Stannard, A.M. Stoker, C.C. Bozynski, K. Kuroki, C.R. Cook, F.M. Pfeiffer, Importance of Donor Chondrocyte Viability for Osteochondral Allografts, Am J Sports Med 44(5) (2016) 1260–8.

[5] J. Parreno, M. Nabavi Niaki, K. Andrejevic, A. Jiang, P.H. Wu, R.A. Kandel, Interplay between cytoskeletal polymerization and the chondrogenic phenotype in chondrocytes passaged in monolayer culture, J Anat 230(2) (2017) 234–248.

[6] F. Mallein-Gerin, R. Garrone, M. van der Rest, Proteoglycan and collagen synthesis are correlated with actin organization in dedifferentiating chondrocytes, Eur J Cell Biol 56(2) (1991) 364–73.

[7] P.D. Benya, J.D. Shaffer, Dedifferentiated chondrocytes reexpress the differentiated collagen phenotype when cultured in agarose gels, Cell 30(1) (1982) 215–24.

[8] A.T. Rzepski, M.M. Schofield, S. Richardson-Solorzano, M.L. Arranguez, A.W. Su, J. Parreno, Targeting the reorganization of F-actin for cell-based implantation cartilage repair therapies, Differentiation 143 (2025) 100847.

[9] J. Abbott, H. Holtzer, The loss of phenotypic traits by differentiated cells. 3. The reversible behavior of chondrocytes in primary cultures, J Cell Biol 28(3) (1966) 473–87.

[10] B. Chan, J. Parreno, M. Glogauer, Y. Wang, R. Kandel, Adseverin, an actin binding protein, regulates articular chondrocyte phenotype, J Tissue Eng Regen Med 13(8) (2019) 1438–1452.

[11] J. Parreno, S. Raju, M.N. Niaki, K. Andrejevic, A. Jiang, E. Delve, R. Kandel, Expression of type I collagen and tenascin C is regulated by actin polymerization through MRTF in dedifferentiated chondrocytes, FEBS Lett 588(20) (2014) 3677–84.

[12] S. Roberts, J. Menage, L.J. Sandell, E.H. Evans, J.B. Richardson, Immunohistochemical study of collagen types I and II and procollagen IIA in human cartilage repair tissue following autologous chondrocyte implantation, Knee 16(5) (2009) 398–404.

[13] B. Kinner, M. Spector, Smooth muscle actin expression by human articular chondrocytes and their contraction of a collagen-glycosaminoglycan matrix in vitro, J Orthop Res 19(2) (2001) 233–41.

[14] M.M. Schofield, A.T. Rzepski, S. Richardson-Solorzano, J. Hammerstedt, S. Shah, C.E. Mirack, M. Herrick, J. Parreno, Targeting F-actin stress fibers to suppress the dedifferentiated phenotype in chondrocytes, Eur J Cell Biol 103(2) (2024) 151424.

[15] J. Parreno, S. Raju, P.H. Wu, R.A. Kandel, MRTF-A signaling regulates the acquisition of the contractile phenotype in dedifferentiated chondrocytes, Matrix Biol 62 (2017) 3–14.

[16] A.L. Pallante, W.C. Bae, A.C. Chen, S. Gortz, W.D. Bugbee, R.L. Sah, Chondrocyte viability is higher after prolonged storage at 37 degrees C than at 4 degrees C for osteochondral grafts, Am J Sports Med 37 Suppl 1(Suppl 1) (2009) 24S–32S.

[17] J.M. Denbeigh, M. Hevesi, C.A. Paggi, Z.T. Resch, L. Bagheri, K. Mara, A. Arani, C. Zhang, A.N. Larson, D.B.F. Saris, A.J. Krych, A.J. van Wijnen, Modernizing Storage Conditions for Fresh Osteochondral Allografts by Optimizing Viability at Physiologic Temperatures and Conditions, Cartilage 13(1_suppl) (2021) 280S–292S.

[18] K.G. Brockbank, E. Rahn, G.J. Wright, Z. Chen, H. Yao, Impact of Hypothermia upon Chondrocyte Viability and Cartilage Matrix Permeability after 1 Month of Refrigerated Storage, Transfus Med Hemother 38(6) (2011) 387–392.

[19] A. Ito, T. Aoyama, H. Iijima, M. Nagai, S. Yamaguchi, J. Tajino, X. Zhang, H. Akiyama, H. Kuroki, Optimum temperature for extracellular matrix production by articular chondrocytes, Int J Hyperthermia 30(2) (2014) 96–101.

[20] A. Ito, T. Aoyama, H. Iijima, J. Tajino, M. Nagai, S. Yamaguchi, X. Zhang, H. Kuroki, Culture temperature affects redifferentiation and cartilaginous extracellular matrix formation in dedifferentiated human chondrocytes, J Orthop Res 33(5) (2015) 633–9.

[21] J. Wang, Y. Wei, S. Zhao, Y. Zhou, W. He, Y. Zhang, W. Deng, The analysis of viability for mammalian cells treated at different temperatures and its application in cell shipment, PLoS One 12(4) (2017) e0176120.

[22] E.E.R. Davis, T.J. Manzoni, V.J. Bianchi, J.F. Weber, P.H. Wu, S.C. Regmi, S.D. Waldman, T.A. Schmidt, A.W. Su, R.A. Kandel, J. Parreno, Passaged Articular Chondrocytes From the Superficial Zone and Deep Zone Can Regain Zone-Specific Properties After Redifferentiation, Am J Sports Med 52(4) (2024) 1075–1087.

[23] J. Parreno, V.J. Bianchi, C. Sermer, S.C. Regmi, D. Backstein, T.A. Schmidt, R.A. Kandel, Adherent agarose mold cultures: An in vitro platform for multi-factorial assessment of passaged chondrocyte redifferentiation, J Orthop Res 36(9) (2018) 2392–2405.

[24] J. Parreno, E. Delve, K. Andrejevic, S. Paez-Parent, P.H. Wu, R. Kandel, Efficient, Low-Cost Nucleofection of Passaged Chondrocytes, Cartilage 7(1) (2016) 82–91.

[25] K.L. Inguito, M.M. Schofield, A.D. Faghri, E.T. Bloom, M. Heino, V.C. West, K.M.M. Ebron, D.M. Elliott, J. Parreno, Stress deprivation of tendon explants or Tpm3.1 inhibition in tendon cells reduces F-actin to promote a tendinosis-like phenotype, Mol Biol Cell 33(14) (2022) ar141.

[26] V.C. West, K.E. Owen, K.L. Inguito, K.M.M. Ebron, T.N. Reiner, C.E. Mirack, C.H. Le, R. de Cassia Marqueti, S. Snipes, R. Mousavizadeh, R.E. King, D.M. Elliott, J. Parreno, Actin Polymerization Status Regulates Tenocyte Homeostasis Through Myocardin-Related Transcription Factor-A, Cytoskeleton (Hoboken) (2024).

[27] R. Mousavizadeh, V.C. West, K.L. Inguito, D.M. Elliott, J. Parreno, The application of mechanical load onto mouse tendons by magnetic restraining represses Mmp-3 expression, BMC Res Notes 16(1) (2023) 127.

[28] M.W. Pfaffl, A new mathematical model for relative quantification in real-time RT-PCR, Nucleic Acids Res 29(9) (2001) e45.

[29] H.J. Motulsky, R.E. Brown, Detecting outliers when fitting data with nonlinear regression - a new method based on robust nonlinear regression and the false discovery rate, BMC Bioinformatics 7 (2006) 123.

[30] V. Ragoonanan, A. Hubel, A. Aksan, Response of the cell membrane-cytoskeleton complex to osmotic and freeze/thaw stresses, Cryobiology 61(3) (2010) 335–44.

[31] M. Krill, N. Early, J.S. Everhart, D.C. Flanigan, Autologous Chondrocyte Implantation (ACI) for Knee Cartilage Defects: A Review of Indications, Technique, and Outcomes, JBJS Rev 6(2) (2018) e5.

[32] J.T. Garrity, A.M. Stoker, H.J. Sims, J.L. Cook, Improved osteochondral allograft preservation using serum-free media at body temperature, Am J Sports Med 40(11) (2012) 2542–8.

[33] R. Shayan-Moghadam, A. Sherafatvaziri, F. Vosoughi, A. Mirzamohamadi, H. Saffar, M. Shafieian, L.O. Zanjani, H. Nematian, M.H. Nabian, Optimum storage conditions for osteochondral allograft plugs: An ex vivo comparative study of 12 storage protocols, J Exp Orthop 12(2) (2025) e70206.

[34] S.M. Frisch, E. Ruoslahti, Integrins and anoikis, Curr Opin Cell Biol 9(5) (1997) 701–6.

[35] H. Arzani, H. Rafii-Tabar, F. Ramezani, The investigation into the effect of the length of RGD peptides and temperature on the interaction with the alphaIIbbeta3 integrin: a molecular dynamic study, J Biomol Struct Dyn 40(20) (2022) 9701–9712.

[36] J. Riegger, M. Zimmermann, H. Joos, T. Kappe, R.E. Brenner, Hypothermia Promotes Cell-Protective and Chondroprotective Effects After Blunt Cartilage Trauma, Am J Sports Med 46(2) (2018) 420–430.

[37] M. Vergara, S. Becerra, J. Berrios, N. Osses, J. Reyes, M. Rodriguez-Moya, R. Gonzalez, C. Altamirano, Differential effect of culture temperature and specific growth rate on CHO cell behavior in chemostat culture, PLoS One 9(4) (2014) e93865.

[38] S. Kaushik, J. Kaur, Effect of chronic cold stress on intestinal epithelial cell proliferation and inflammation in rats, Stress 8(3) (2005) 191–7.

[39] I.C. Enninga, R.T. Groenendijk, A.A. van Zeeland, J.W. Simons, Use of low temperature for growth arrest and synchronization of human diploid fibroblasts, Mutat Res 130(5) (1984) 343–52.

[40] N.C. Leegwater, A.D. Bakker, J.M. Hogervorst, P.A. Nolte, J. Klein-Nulend, Hypothermia reduces VEGF-165 expression, but not osteogenic differentiation of human adipose stem cells under hypoxia, PLoS One 12(2) (2017) e0171492.

[41] R.J. Williams, 3rd, J.C. Dreese, C.T. Chen, Chondrocyte survival and material properties of hypothermically stored cartilage: an evaluation of tissue used for osteochondral allograft transplantation, Am J Sports Med 32(1) (2004) 132–9.

[42] C. Albrecht, B. Tichy, S. Nurnberger, L. Zak, M.J. Handl, S. Marlovits, S. Aldrian, Influence of cryopreservation, cultivation time and patient’s age on gene expression in Hyalograft(R) C cartilage transplants, Int Orthop 37(11) (2013) 2297–303.

[43] A. von Bomhard, J. Faust, A.F. Elsaesser, S. Schwarz, K. Pippich, N. Rotter, Impact of expansion and redifferentiation under hypothermia on chondrogenic capacity of cultured human septal chondrocytes, J Tissue Eng 8 (2017) 2041731417732655.

[44] D.Y. Kim, K.M. Kim, E.J. Kim, W.G. Jang, Hypothermia-induced RNA-binding motif protein 3 (RBM3) stimulates osteoblast differentiation via the ERK signaling pathway, Biochem Biophys Res Commun 498(3) (2018) 459–465.

[45] Y. Zhu, H. Tao, C. Jin, Y. Liu, X. Lu, X. Hu, X. Wang, Transforming growth factor-beta1 induces type II collagen and aggrecan expression via activation of extracellular signal-regulated kinase 1/2 and Smad2/3 signaling pathways, Mol Med Rep 12(4) (2015) 5573–9.

[46] J.G. Edwards, J.A. Campbell, The aggregation of trypsinized BHK21 cells, J Cell Sci 8(1) (1971) 53–71.

[47] B.M. Gumbiner, Cell adhesion: the molecular basis of tissue architecture and morphogenesis, Cell 84(3) (1996) 345–57.

[48] E. Ruoslahti, Integrins as signaling molecules and targets for tumor therapy, Kidney Int 51(5) (1997) 1413–7.

[49] D.J. Andlinger, U. Kulozik, Protein-protein interactions explain the temperature-dependent viscoelastic changes occurring in colloidal protein gels, Soft Matter 19(6) (2023) 1144–1151.

